# Dendritic delay lines shape the computation of sound location in neurons of the gerbil medial superior olive

**DOI:** 10.64898/2026.01.28.702328

**Authors:** Jared Casarez, Rebecca I. Voglewede, Bradley D. Winters, Ken Ledford, Nace L. Golding

**Author notes:** Equal contributions (Jared Casarez, Rebecca I. Voglewede). **Corresponding author and lead contact** Nace L. Golding: The University of Texas at Austin, Department of Neuroscience and Center for Learning and Memory, 1 University Station C7000, Austin TX 78712-0248,.

## Abstract

In mammals, neurons of the medial superior olive (MSO) encode information about sounds arising from discrete spatial locations along the horizon. This tuning requires that an internal delay in the brain must offset acoustic disparities to ensure coincident arrival of excitatory inputs driven from the two ears. The source of this optimal internal delay, originally assumed to arise from axonal delay lines, is currently controversial in mammals. Here we use 2-photon guided paired dendritic and somatic recordings together with compartmental modeling of 40 complete MSO neuron morphologies to demonstrate that the dendrites themselves serve as a significant source of internal delay. We show that most MSO neurons exhibit morphological asymmetries that impose different EPSP delays across dendrites and shifts in optimal interaural time differences. Dendrite-based delays in the mammalian MSO are heterogeneous within each isofrequency laminae and provide a stable, structural mechanism to help tune individual neurons to sounds from different azimuthal locations.

## INTRODUCTION

Spatial information in the auditory system is critical for humans and other mammals to assemble a three-dimensional view of the world as well as to understand speech and communication signals, especially in environments with many sounds overlapping in time and in frequency. The auditory system is distinct from vision or touch in that there is no explicit peripheral representation of space, in this case in the cochlea, only a representation of frequency. Accordingly, the central auditory system must compute horizontal sound location indirectly by combining a series of monaural and binaural cues (Grothe and Pecka, 2014; Joris and Yin, 2007). Spatial coding in the central auditory system is a prime example of a centrally synthesized or computational map.

Interaural time difference cues are used by birds and mammals to compute horizontal sound location at low frequencies (Carr et al., 2001). In mammals this computation takes place in the medial superior olive (MSO), whose neurons are the first to receive converging excitatory and inhibitory inputs driven from the two ears. Intriguingly, in individual neurons, binaural inputs are integrated within a bipolar dendritic architecture, with ipsilateral and contralateral excitatory inputs segregated onto lateral and medial dendritic branches respectively (Lindsey, 1975; Stotler, 1953). Ipsilateral and contralateral inhibition converges onto the soma and proximal dendrites (Kapfer et al., 2002; Lindsey, 1975; Magnusson et al., 2005; Stotler, 1953; Yin and Chan, 1990). MSO neurons detect the optimal combination of these inputs, converting temporal coincidence into a rate code (Goldberg and Brown, 1969; Spitzer and Semple, 1995; Yin and Chan, 1990).

The concept of internal delay is central to coincidence detection in the MSO, where acoustic delays to the two ears are countered in time by a mechanism of delay inside the brain. This internal temporal asymmetry was postulated in early models to arise from axonal delay lines formed by ladder-like combinations of bilateral excitatory inputs (the “Jeffress model”) (Jeffress, 1948). While the Jeffress model enjoys strong support in birds (Carr and Konishi, 1990, 1988; Moiseff and Konishi, 1981), comparable studies in mammals have failed to reveal systematic axonal delay lines nor any mapping of delay along the nucleus (Beckius et al., 1999; Smith et al., 1991; Yin and Chan, 1990). More recently, there is evidence that fast inhibition mediates internal delay by shifting the peaks of EPSPs (Brand et al., 2002; Myoga et al., 2014; Pecka et al., 2008). However, this mechanism is subject to vigorous debate (Franken et al., 2015; Heijden et al., 2013; Roberts et al., 2013), and it is not clear that this mechanism alone can explain the range of receptive field locations in the MSO or their insensitivity to sound level. Other mechanisms have been proposed to mediate internal binaural asymmetry, including asymmetrical axon position, medio-lateral differences in the rising slopes of EPSPs (Jercog et al., 2010), and differences in tuning of binaural inputs (“stereausis”) (Plauška et al., 2017; Shamma et al., 1989).

All models of coincidence detection must address the issue of the temporal distortions imposed by the dendrites themselves. However, to date models of MSO neurons have been represented either as point neurons or as simplified, symmetrical “ball-and-stick” models (Agmon-Snir et al., 1998; Callan et al., 2021; Dasika et al., 2007; Franken et al., 2014; Mathews et al., 2010; Myoga et al., 2014). Previous measurements of dendritic propagation have revealed that EPSPs encounter delays in their propagation to the soma up to ∼150 µs, despite some degree of temporal compensation by activation of low voltage-activated K channels in the soma and proximal dendrites (Mathews et al., 2010). However, these measurements have largely been restricted to the proximal half of the dendritic arbor and have come from averaging highly variable populations of neurons. Given that the dendrites of MSO neurons alter their branching pattern and diameters during early hearing and following disruptions of normal hearing (Feng and Rogowski, 1980; Russell and Moore, 1999; Smith and Rubel, 1979), they might serve as a potential mechanism for adjusting the timing requirements for sounds from the two ears to arrive coincidently at the soma and axon.

Here we demonstrate that the dendritic arbors of MSO neurons are highly heterogeneous and show a striking continuum of asymmetry across the medial and lateral branches that receive segregated inputs driven respectively from the two ears. We show with dual 2-photon fluorescence-guided patch recordings that inputs to smaller and/or more distal dendrites show more substantial propagation delays than previously recognized, and computational modeling of a large population of intact neuronal morphologies shows that these dendritic asymmetries translate into stable internal delays that, as a population, could account for the full range of naturally occurring horizontal locations. Finally, we show that inhibition preserves the effects of dendritic asymmetry on interaural time differences (ITDs) and helps bring the region of maximum slope of firing changes within the ecologically relevant range. Thus dendrite-based delay lines likely work synergistically with other integrative mechanisms purported to shape the perception of horizontal position in mammals.

## RESULTS

To understand how MSO neurons process interaural time differences in the dendrites, we made fluorescence-guided dendritic and somatic patch-clamp recordings visualized under 2-photon microscopy from MSO neurons in gerbil brainstem slices near physiological temperature (35°C). Using this approach, we were able to target the distal third of the dendritic arbor, including secondary and tertiary branches, regions that have largely escaped investigation in these neurons. We then injected simulated EPSCs (τ_rise_ and τ_decay_, 0.2 ms) into the dendritic pipette and recorded potentials simultaneously at both dendrites and soma. We observed strong variability in both the percent amplitude attenuation at the soma as well as the propagation delay of the EPSP, measured as the difference in EPSP peak time at the soma relative to the dendrite (Fig. 1). Notably, the delays measured in these recordings ranged between 100 and 300 µs, exceeding the physiological range of ITDs measured in gerbils. In these recordings, there was little correlation between local morphological features of recorded dendrites and propagation delay of the EPSP, measured as the difference in the peak timing of dendritic and somatic EPSPs (Fig. 2a-c). However, there was a weak correlation between the local dendritic membrane time constant and EPSP propagation delay (Fig. 2d). In addition, there was a weak trend of increasing dendritic membrane time constant with recording distance (Fig. 2e), possibly because of proximity to the dendritic terminations. These results suggest that an understanding of the integrative properties of the dendrites requires an analysis of the larger morphological and electrotonic structure of the neuron.

**Fig. 1:**
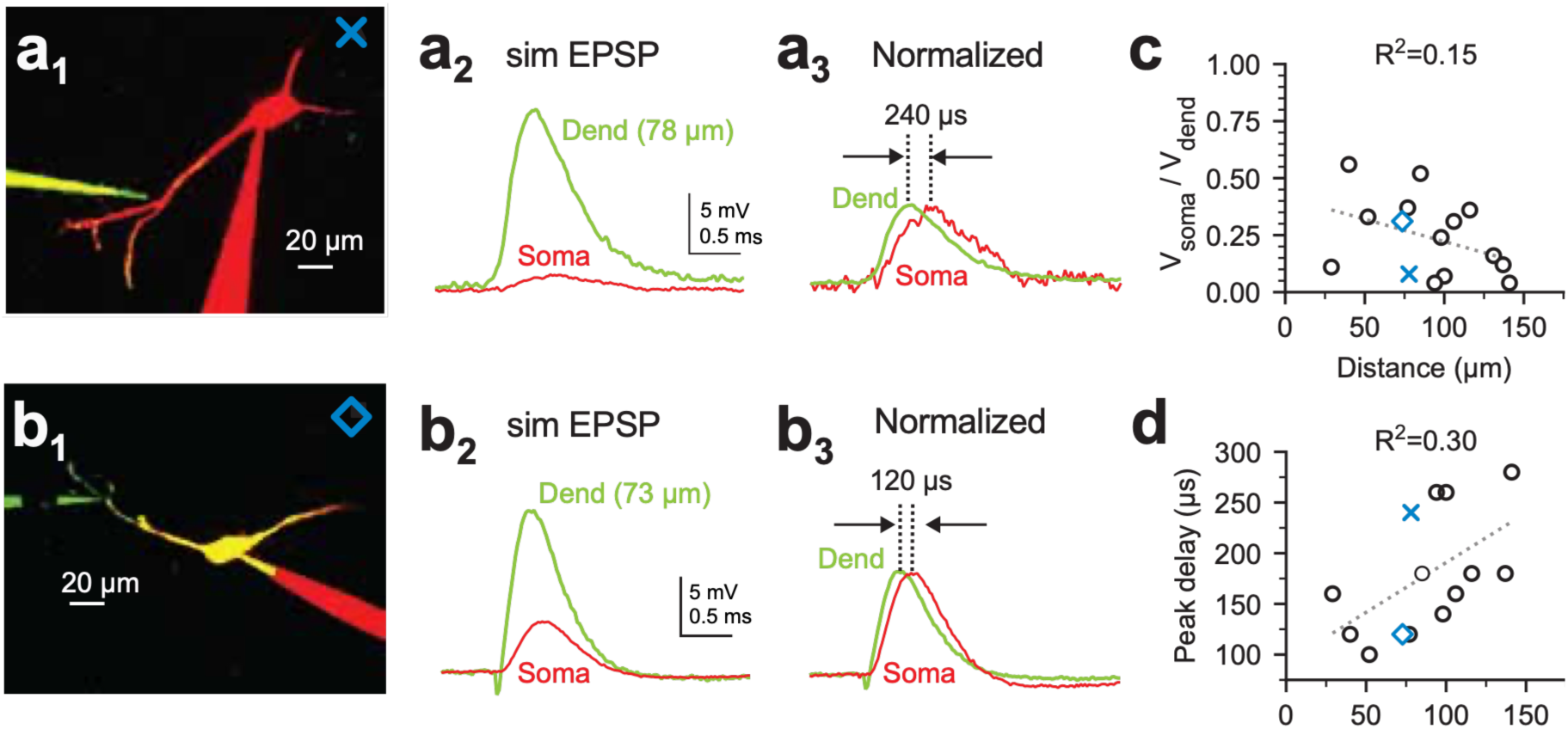
Heterogeneity in attenuation and timing of simulated excitatory synaptic potentials following propagation from distal dendritic locations to the soma. a,b, 2-photon fluorescence-guided dual dendritic and somatic patch recordings from two MSO neurons of comparable distances (78 and 73 µm, respectively). Example neuron in **a1** shows stronger attenuation of somatic propagation of a simulated EPSP (**a2** vs. **b2**) and longer peak delay (**a3** vs. **b3**). EPSC amplitudes: cell **a**, 600 pA; cell **b**, 400 pA. **c,d**, Group data for attenuation and peak delay for all paired MSO neuron recordings. There is a poor correlation of both measures as a function of distance.

**Fig. 2:**
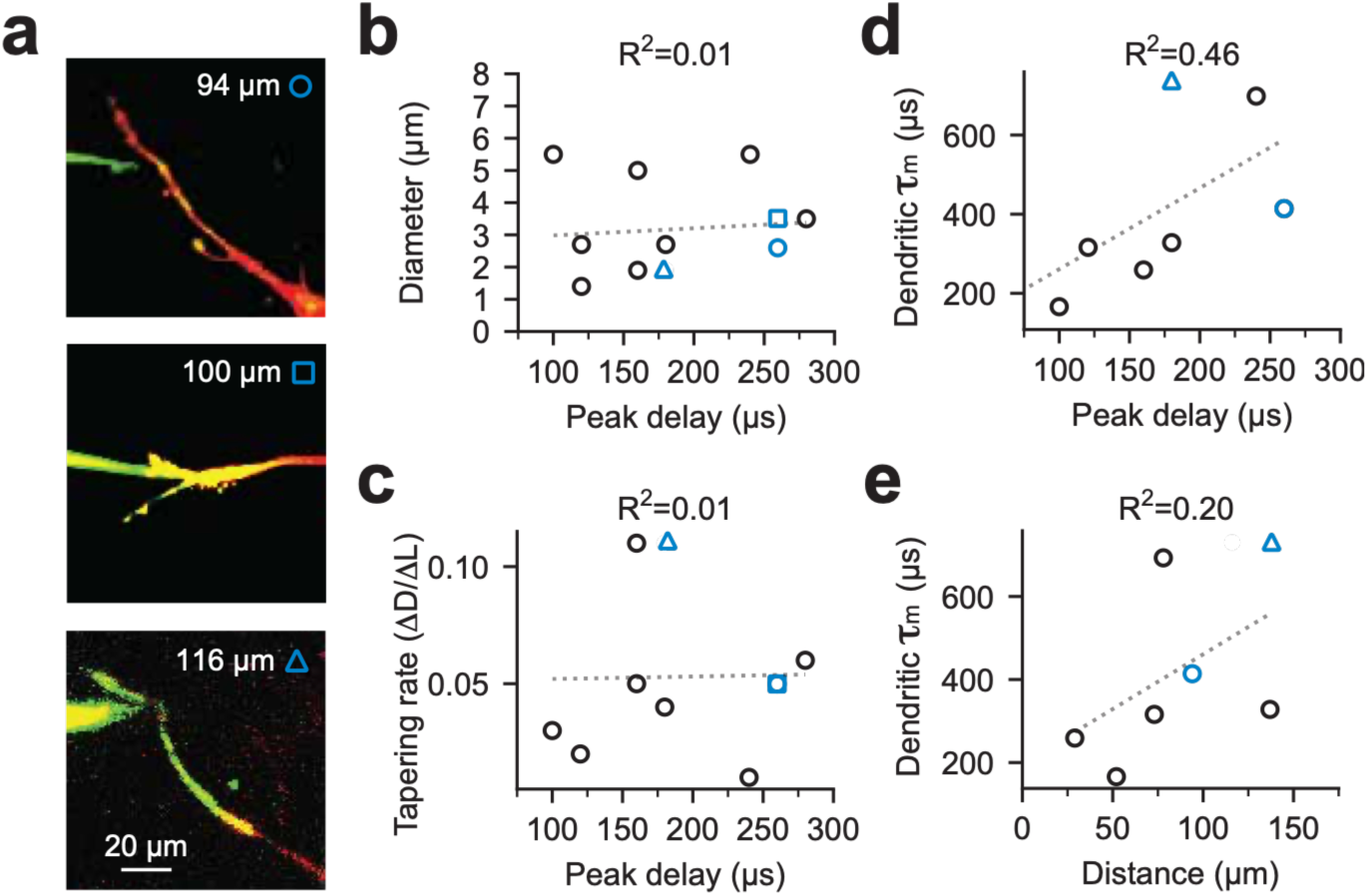
EPSP propagation delays are not correlated with local morphology but rather more global physiological properties. **a**, Closeups of distal dendritic morphology during paired dendritic and somatic recordings. **b,c**, Delay of peak EPSPs during propagation from dendrite to soma shows no significant correlation with local diameter or tapering rate of local dendrite. (dendritic EPSCs: 400-1000 pA). **d**, Peak EPSP delay from dendrite to soma was weakly correlated with the membrane time constant recorded at the dendritic recording site. **e**, Dendritic membrane time constant showed only a weak positive trend with distance of the recording site from the soma.

Given that the EPSP propagation delays from the distal dendrites to the soma are substantial compared to the ecologically relevant range of ITDs available to the gerbil, we hypothesized that differences in length, diameter or branching structure of the medial and lateral dendrites of individual cells could significantly impact the temporal requirements for EPSP summation and binaural coincidence detection. To pursue this question, we analyzed a database of biocytin-filled MSO neurons that exhibited complete dendritic structures (40 of 270 neurons from Bondy et al., 2021; Fig. 3). We quantified dendritic length, surface area and branching, with special emphasis on comparisons of medial and lateral dendritic arbors. Higher-order (2° and higher) dendritic branches were prominent in these neurons, comprising 70% of the membrane surface area (Figs. S1 and S2).

**Fig. 3:**
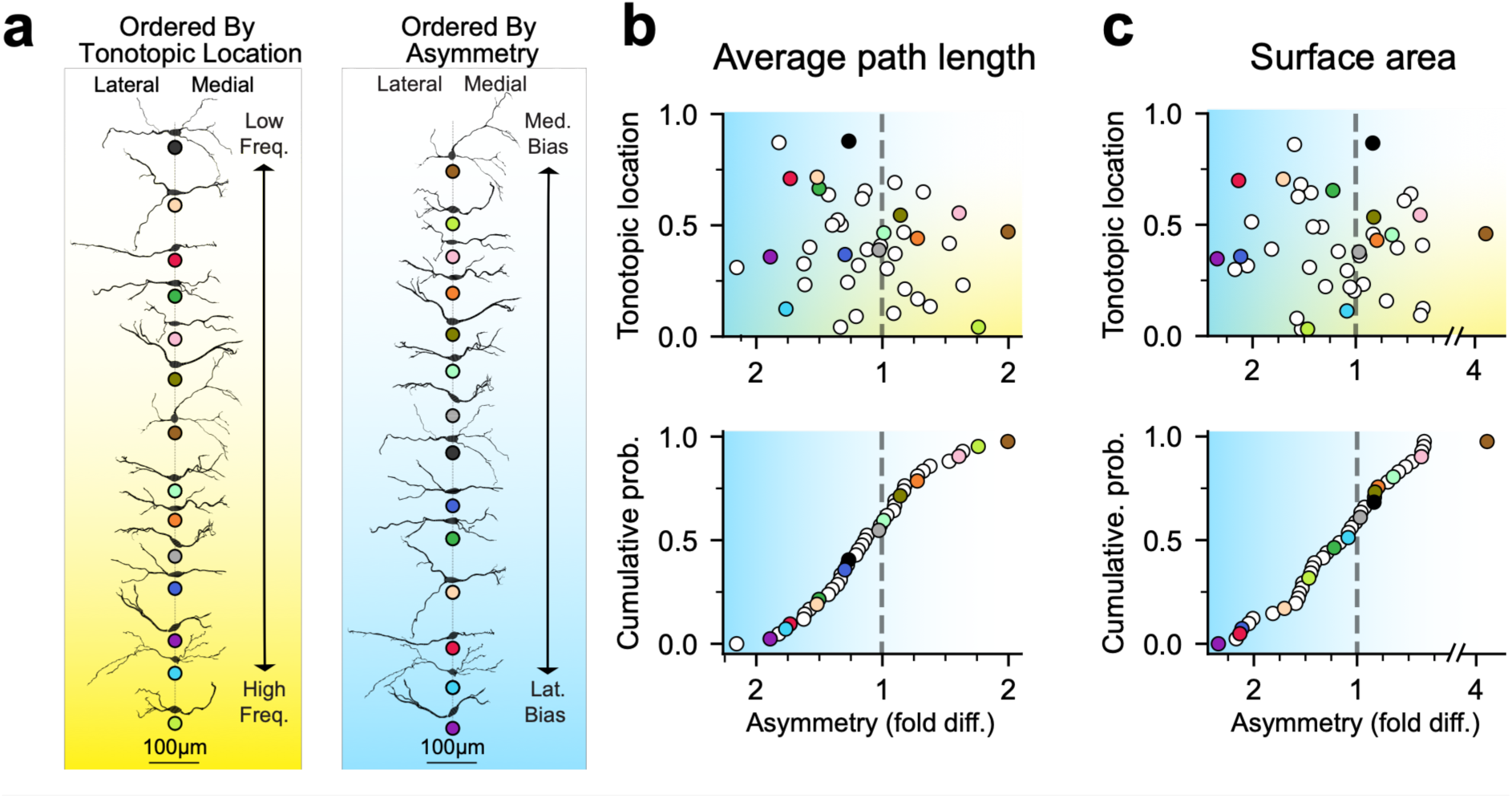
MSO neuron morphology is inhomogeneous and asymmetric, regardless of tonotopy. **a**, Reconstructed morphologies rank ordered by tonotopy (left) and degree of asymmetry (right; average path length). Orderings span from low frequency (top) to high frequency (bottom) and medial (top) and lateral (bottom) asymmetry, respectively. Colored gradients added to visualize ordering. Uniquely colored dots placed beneath each morphology for differentiation. **b**, Fold difference between average path length of medial and lateral dendrites for all reconstructed morphologies (n=40) with vertical axes ordered by tonotopic location (top) or degree of asymmetry (bottom). Gradients added as in **a**. Colored points correspond to **a**. **c**, Same as **b**, except comparing total surface area of medial and lateral branches. Average medio-lateral path length and surface area fold differences (± SEM) were 1.080 ± 0.057 and 1.112 ± 0.133 (biased laterally) and tonotopic R^2^ values were -0.42 and -0.23.

A striking feature of most neurons in this data set was an overall asymmetry between the lateral and medial dendritic arbors, reflecting differences in length, branching, and/or diameter of the dendrites (Fig. 3a-c). We quantified this asymmetry according to two parameters, the average path length (average length from each dendritic end to the soma) and membrane surface area, with the latter parameter reflecting the collective influence of dendritic length, diameter and branching pattern. We then defined the degree of bilateral asymmetry as fold differences between the larger vs. smaller dendritic measure for each parameter. We observed no obvious correlation between dendritic asymmetry and tonotopic location (Fig. 3a, *left* and 3b, *top*). However, we found that dendritic asymmetry formed a finely graded continuum, with a slight bias toward lateral dendrites (Fig. 3b, *bottom*). Interestingly, there were no significant differences in population averages of the medio-lateral path length and surface area (1.080 ± 0.057 and 1.112 ± 0.133-fold differences; p=0.562), though there was a slight lateral bias. These results indicate that the averaging of dendritic morphological parameters across neurons obscures what is likely a key structural and computational feature.

To understand how morphological asymmetry impacts synaptic integration, we constructed compartmental models using the same 40 reconstructions analyzed in Fig. 3. We introduced an excitatory synaptic conductance (alpha function: t = 290 µs) into each dendritic compartment and then measured the voltage attenuation and propagation delay from the dendritic input location to the soma (Figs. 4–5). EPSP amplitude varied considerably both at the site of synaptic input and soma despite comparable distances, like the physiological data from paired recordings in Figure 1 (Fig. 4a). Plots of local dendritic and propagated somatic EPSP amplitudes across all synapse locations further revealed striking asymmetries in the attenuation of EPSPs to the soma across specific branches even when arising from comparable distal locations (Fig. 4b, “Active cell”, red traces in the leftward oriented (lateral) dendrites distal to ∼50 µm). In MSO neurons, the timing and attenuation of EPSPs along dendrites are influenced both by passive cable filtering as well as interactions with low voltage-activated K channels, which are active in the subthreshold voltage range in these neurons (Mathews et al., 2010). To determine the degree to which asymmetries in low voltage-activated K channels impact EPSP attenuation at the soma we converted all voltage-activated conductances to leak channels (Fig. 4b, “Passive cell”; see Methods). While EPSPs propagated slightly more effectively to the soma from all dendritic locations in the passive model, it did not fundamentally disrupt mediolateral asymmetry, indicating that these differences arose primarily from the passive electrotonic architecture of the cell. The branching structure had a significant effect on dendritic propagation asymmetry. This could be better visualized in morphoelectrotonic transforms of cells, which display dendritic compartments according to their electrotonic, versus physical lengths. In the example neuron in Fig. 4c, large primary dendrites are deemphasized, while thinner secondary and tertiary branches appear as electrotonically elongated. Differences between the medial and lateral dendritic arbors in this neuron and many others gave rise to asymmetries in the magnitude of the average EPSP arriving at the soma.

**Fig. 4:**
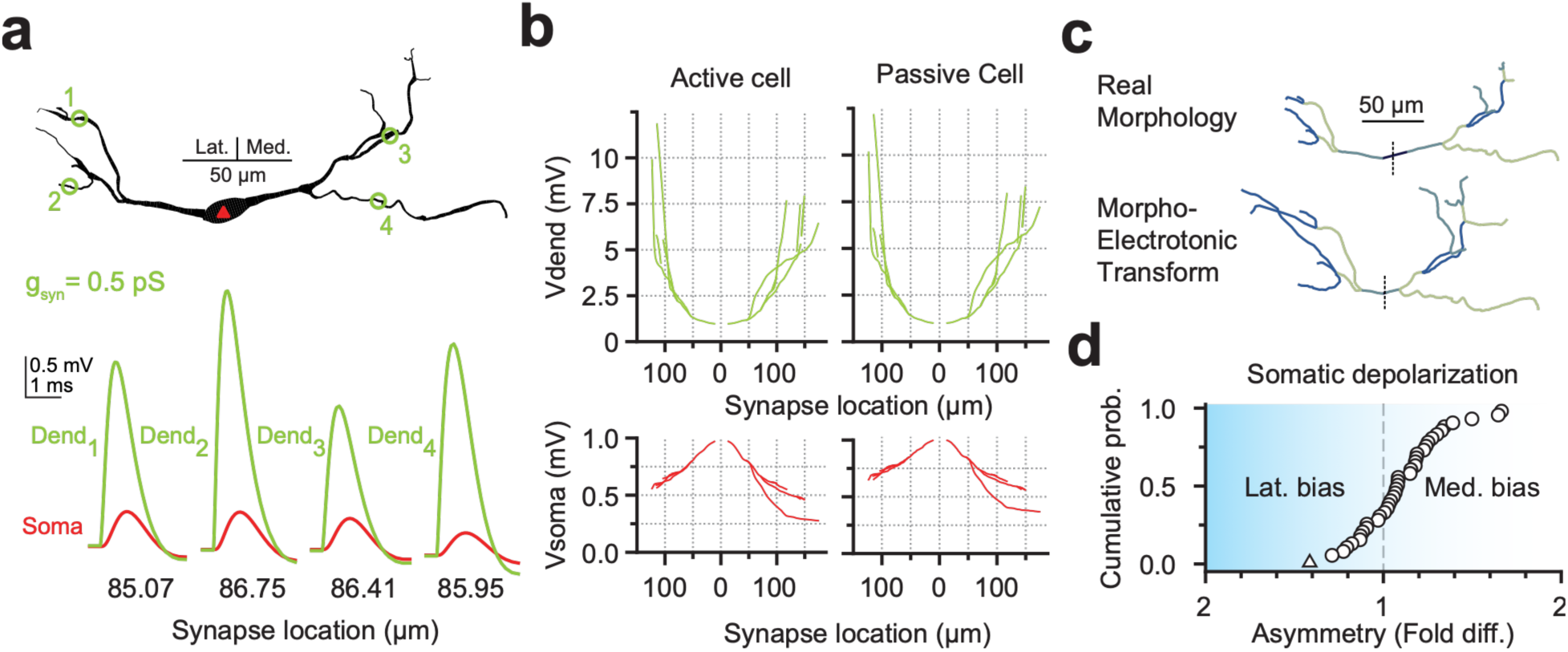
MSO morphology nonlinearly filters and attenuates propagating EPSPs through its passive cable properties. a, Visualization of EPSP filtering at different dendritic sites. Cell morphology (top) with similarly distant example synaptic sites indicated with green circles and corresponding numbers. Red triangle indicates the somatic compartment. Synaptic conductance indicated as gsyn. Four sets of paired dendritic (green) and somatic (red) traces (bottom) for each example synaptic site. Numbering of traces corresponds to labels on morphology above. Dendritic distance from soma for each site indicated below traces. **b**: Same procedure as in **a**, with an adjusted unitary synaptic conductance (see Methods), with measurements from each computational compartment. Model morphology and orientation same as in **a**. Comparison between active channel (left) and passive channel model (right). Recorded EPSP amplitude at synaptic (top, green) and somatic (bottom, red) compartment as a function of distance from soma. **c**, Actual (top) and electrotonically transformed (bottom) morphologies. Transformation visualizes degree of EPSP attenuation from each compartment (see Methods). **d**, Group data (n=40) of average somatic depolarization asymmetry between mediolateral sides. Gradient as in Fig 3. Dotted line indicates equal medial and lateral average somatic depolarizations. Cell from **a-c** indicated with a triangle.

**Fig. 5:**
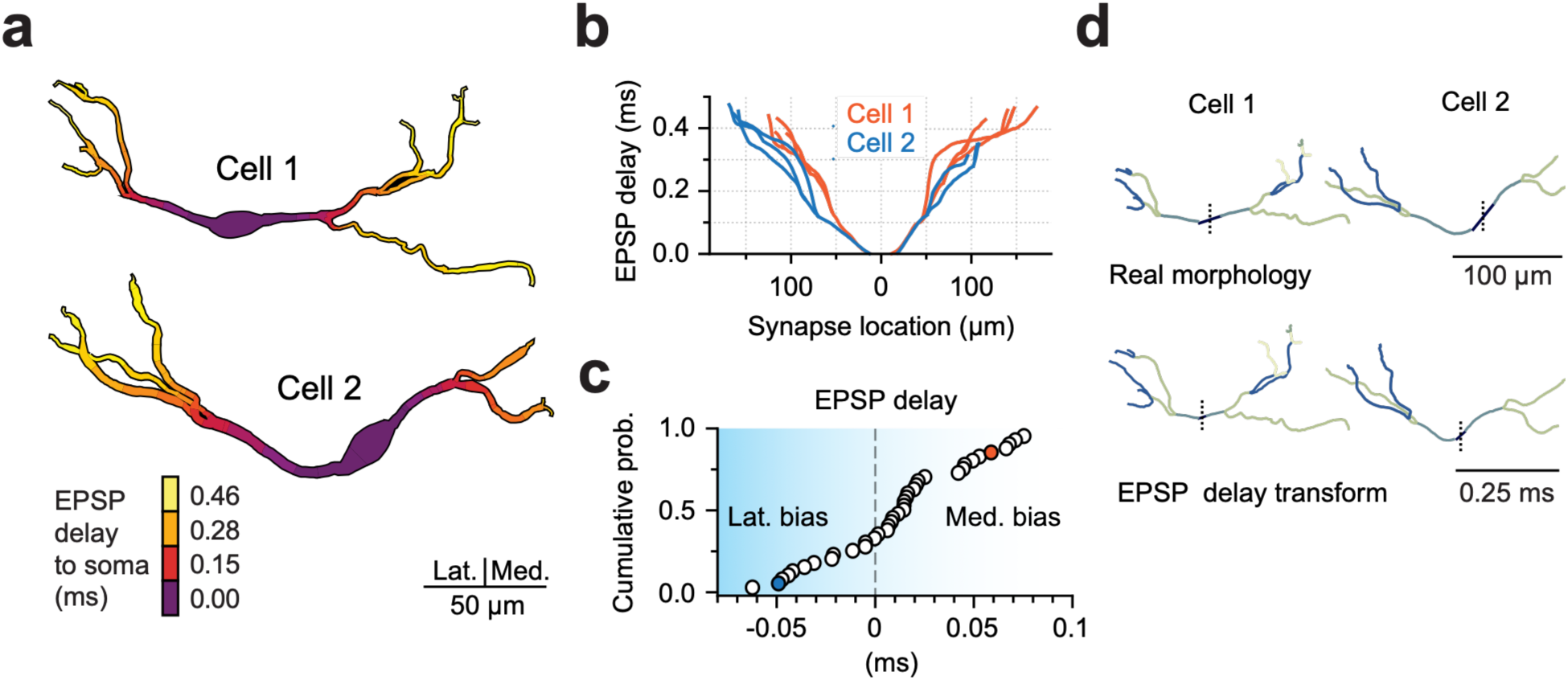
EPSP timing significantly varies within and between cells, with morphological asymmetry informing overall cell timing. **a,** EPSP delay visualization in two exemplar MSO neurons. Coloring of each cell corresponds to the propagation delay of EPSPs to the soma from its respective compartment. Legend of propagation delay coloring in bottom left. Cells are oriented mediolaterally according to the scale bar. Outline added to increase gradient readability. **b,** Same data as in **a**, except as a function of distance from the soma. Orientation of cells same as in **a. c**, Group data (n=40) of the difference in average EPSP propagation delay between medial and lateral dendrites. Cells 1 & 2 are indicated with same color as in **b**. Dotted line at 0 ms indicates an equal average EPSP delay between sides. Blue gradient along horizontal axis (as in Fig. 3) to indicate spectrum of asymmetry. **d**, Morphological transforms according to EPSP delays. Centroids of actual cell morphologies (top) and transformed morphologies (bottom). Length corresponds to each section’s EPSP delay contribution. Coloring of cell sections is consistent between actual and transformed morphologies for easier differentiation and comparison. Dotted line: soma midline.

Further, we observed that our population of cells formed a continuum in the degree of somatic depolarization asymmetry, in line with the asymmetry in dendritic structure (Fig. 4d). Dendritic asymmetry also translated into differences in average EPSP timing across the medial and lateral dendrites. Figure 5a shows for two example neurons the spatial profile of delays of EPSPs propagating from each dendritic compartment to the soma. The asymmetry in dendritic timing can be more quantitatively visualized in plots of peak EPSP delay (t_dend_ – t_soma_) as a function of dendritic distance from the soma (Fig. 5b). Dendritic branch points often introduced substantial jumps in propagation delays, and this is reflected in these plots as increases in slope. To quantify the influence of dendritic asymmetry on the optimal summation of bilateral EPSPs at the soma, we measured the electrotonic delay at the soma for every 2 µm compartment of lateral and medial dendritic subtrees. In our population of 40 neuron models, there were differences in the average relative time of arrival of EPSPs to the soma from compartments in the medial and lateral dendrites. Further, these differences formed a smoothly graded continuum of delays (Fig. 5c). These results can be visualized in the morphoelectrotonic transforms for delay for an example neuron (Fig. 5d).

MSO neurons exhibit a range of intrinsic electrical properties, and so we examined a subset of neuron morphologies to understand the functional implications of this physiological diversity. Using a neuron with an asymmetric dendritic arbor, we systematically slowed the membrane time constant by reducing the density of both low voltage-activated K channels and HCN channels (Fig. S3). In both cases, the timing of EPSPs reaching the soma was linearly related to the value of the membrane time constant. Thus, the effects of dendritic asymmetry on EPSP timing and amplitude at the soma are expected to be substantially higher in MSO neurons with slower intrinsic membrane properties. To understand how the differential timing of EPSPs propagating from medial and lateral dendrites is translated into ITDs, we simulated binaural excitatory input using a spiking model of MSO neurons (see Methods). This model featured one axonal input per terminal branch segment as well as a centripetal (dendrite-to-soma) pattern of dendritic innervation, running from the terminal compartments to the soma, in accordance with results from anatomical studies (Fig. 6a) (Beckius et al., 1999; Callan et al., 2021; Smith et al., 1991). The conduction velocity of these inputs was set to 1 m/s, approximating values of unmyelinated terminal axonal arborizations of similar caliber (Carr and Konishi, 1990; Hoffmeister et al., 1991). The timing of ipsilateral and contralateral inputs was varied systematically, triggering spiking only when the two inputs were in close coincidence (Fig. 6b). The location of these excitatory ITD functions were dependent on medio-lateral asymmetry. Functions shifted toward the side of the electrically longer dendrite, reflecting the need for earlier activation of that side to compensate for a longer dendritic propagation delay (Fig. 6c). Our population of 40 neurons showed peak firing at delays that traversed a continuum of values ( -0.11 to 0.08 ms, median -0.02; Fig. 6d; Fig. S4). The precise range of values depended in part on the assumptions on the velocity of excitatory axons. The largest range of ITDs was achieved when there was minimal difference between the velocity of axonal input velocity and postsynaptic EPSPs propagating from the dendrites to the soma, which maximized temporal coincidence at the soma of spatially disparate inputs in the dendrites (Fig. 6e). Finally, ITDs shifted less than 3% in the face of input amplitude changes that resulted in firing probabilities between 0.2 and 0.8, indicating that the influence exerted by dendritic asymmetry on receptive field position is highly stable across changing stimulus conditions (Fig. 6f).

**Fig. 6:**
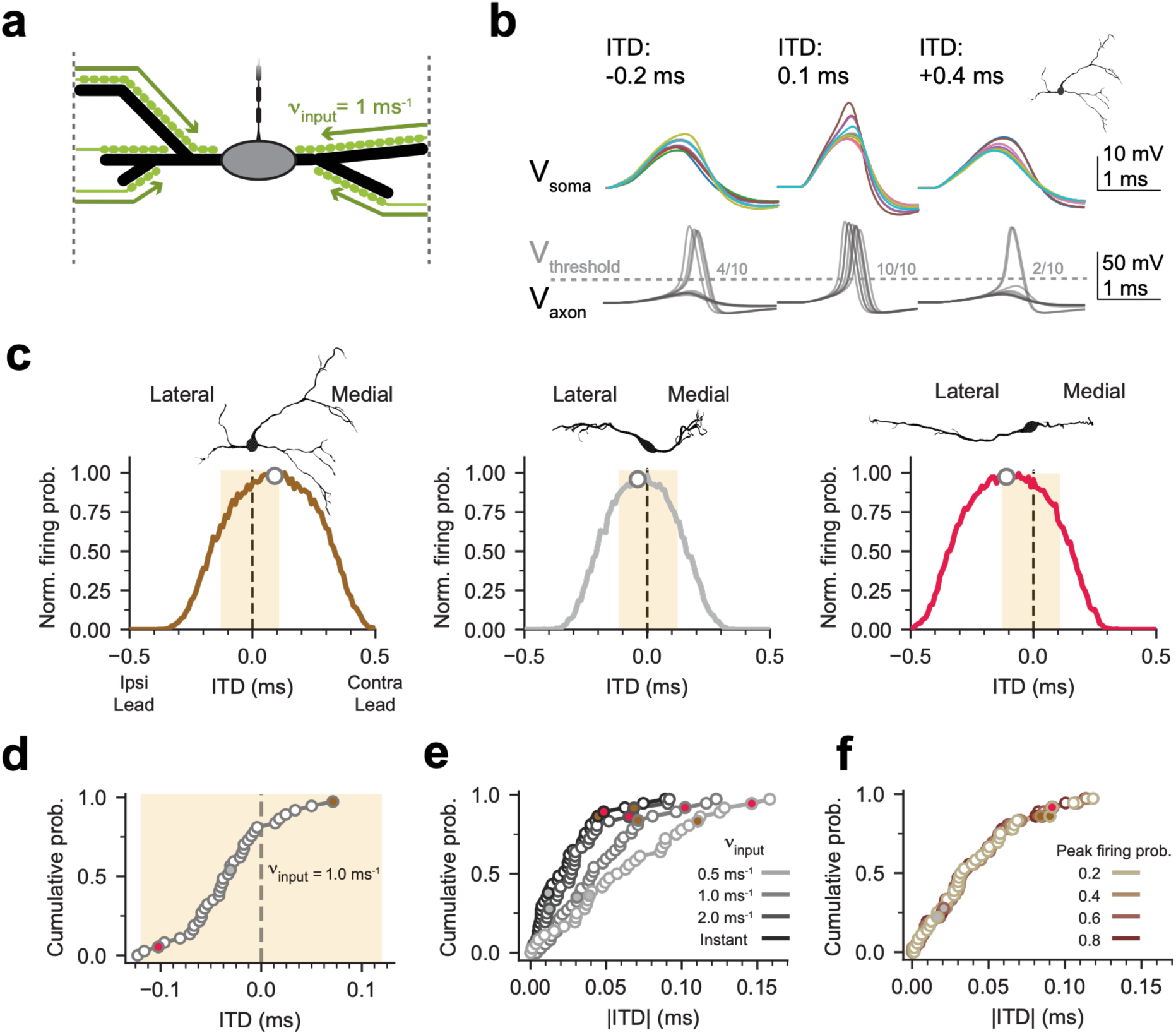
Simulated ITDs demonstrate conservation of morphological impact on coincident detection. **a**, Schematic visualization of ITD innervation. Somatic compartment and dendrites are shown in gray and black, respectively. Green bulbs represent excitatory synapses, with corresponding arrows to illustrate axonal input velocity (νinput). Dotted lines indicate the distal locations where t=0 (see Methods). Axon attached to soma, with initial segment, nodes, and internodes. **b**, Example somatic (top) and axonal (bottom) traces from ITD simulations. ITDs of -0.2 ms (left), 0.1 ms (middle, optimal coincidence), 0.3 ms (right) shown. Traces colored only for differentiation. Dotted line (Vthreshold) indicates depolarization threshold for action potential consideration. Morphology of model cell indicated in upper right. **c**, Three example ITD firing curves from varied morphologies, medially biased (left), symmetrical (middle), and laterally biased (right). Light orange box represents range of ecologically possible ITDs for model species (±130 µs). Dashed line placed at ITD of 0 ms. Dot indicates centroid ITD value. **d,** Sorted group data (n=39) of centroid ITDs, with νinput = 1 m/s. Cells from **c** indicated with filled colored points. **e,f**, Sorted group data (n=39) of absolute centroid ITDs, varying axonal input velocity (**e**) and peak firing probability (**f**). Cells from **c** indicated with correspondingly filled colored points.

Inhibition is a critical component of ITD encoding, although its precise role is subject to vigorous debate. To understand how dendritic asymmetry impacts the effects of inhibition, we added bilateral inhibitory conductances to the somatic compartment of the bilateral excitatory model in Fig. 6, consistent with the known somatically biased innervation of MSO neurons from the lateral and medial nuclei of the trapezoid body (LNTB and MNTB; (Fig. 7a; Kapfer et al., 2002; Werthat et al., 2008)). We set the timing of inhibitory conductances to be 500 µs in advance of excitatory inputs from the same side, in accordance with a previously published estimate (Roberts et al., 2013). We then adjusted the amplitude of inhibitory conductances to produce summed IPSPs of between ∼1-3 mV, which reduced firing probability up to ∼80% (Fig. 7b, c). With increasing amplitude of

**Fig. 7:**
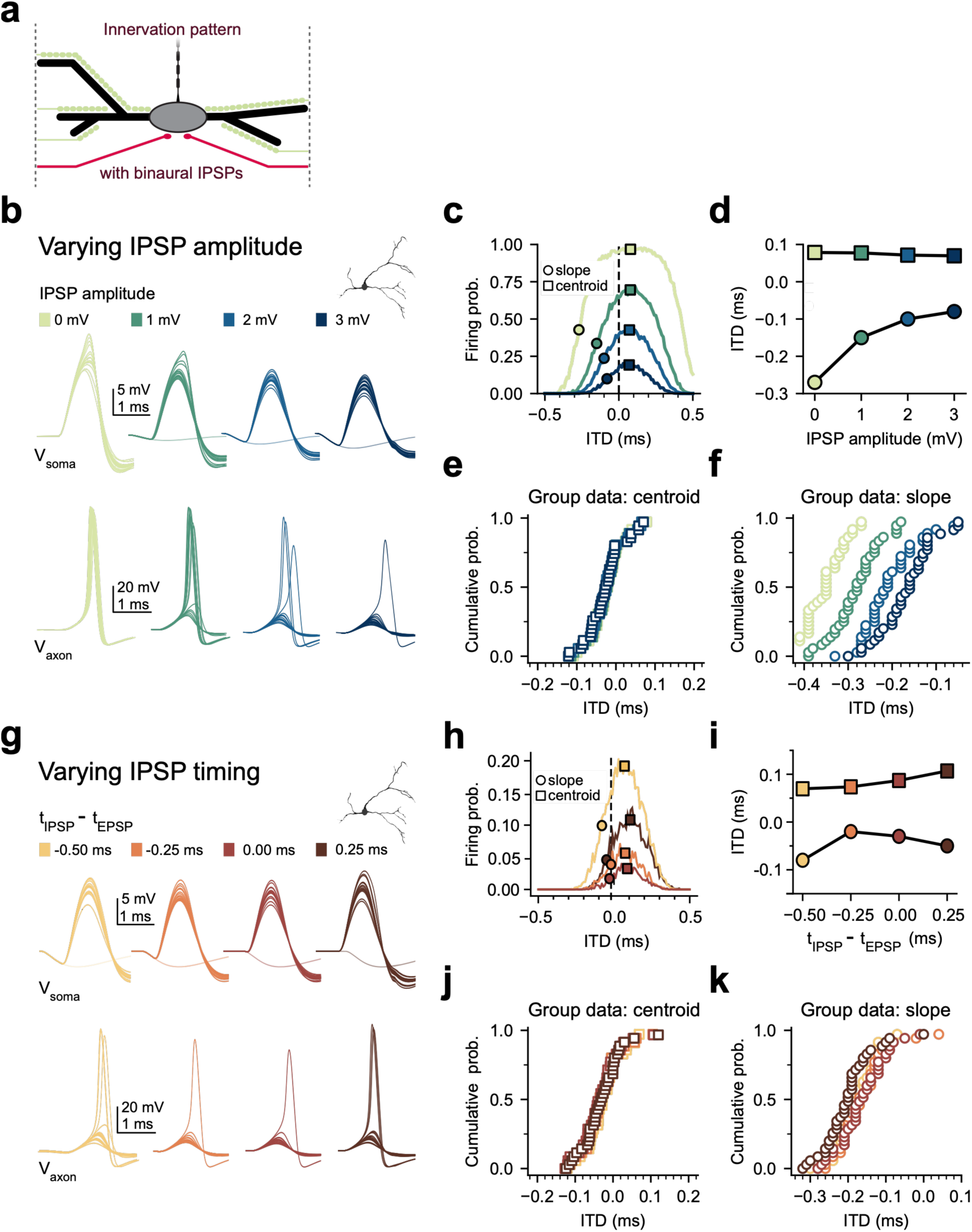
Inhibition may serve to modulate a pre-established morphological baseline ITD response. a, Schematic of innervation pattern with excitatory (green) and inhibitory (red) inputs. **b,** Somatic (top) and axonal (bottom) traces from ITD stimulation (ITD: 0.1 ms; IPSP timing: -0.5 ms) for each tested IPSP amplitude. Tested amplitude values and their unique colors above corresponding traces. Morphology of the exemplar neuron shown in the upper right. **c,** Exemplar cell’s ITD response curves colored corresponding to IPSP amplitude in **b.** Centroid (square) and maximum slope (circle) are indicated for each curve. Dotted line: ITD = 0 ms. **d,** ITD centroid and maximum slope as a function of IPSP amplitude (colors corresponding to **b**). **e,f**, Group data (n=39) for ITD centroids (**e**) and maximum slope (**f**) for each IPSP amplitude. Data in **e** are largely overlapping. Shifting of population data in **f** corresponds to trend seen in the exemplar cell (**d**). **g-k**, Same as **b-f**, instead varying the timing of the inhibition (IPSP amplitude: 3 mV). Absolute firing probability, in **h**, is lower than **c**, resulting in lower signal to noise ratio. Centroids in **i** show more variation than **d**, though still minimal. Shift of exemplar cell’s maximum slope ITD section (**i**) corresponds to degree of reduction in absolute firing probability (**h**).

IPSPs, resultant ITD functions decreased in halfwidth by 57%, but the location of functions with increasing levels of inhibition were not significantly different from the model with only excitatory inputs (0.070-0.078 ms; centroid, p=0.984; group data, Fig. 7d, squares). However, the region of maximum slope consistently shifted towards zero, in accordance with a sinking iceberg receptive field scenario (Fig. 7d, circles; -0.27 to 0.07 ms). We then tested how the relative timing of excitatory and inhibitory inputs altered ITD functions. Varying the timing of IPSPs relative to EPSPs from –0.5 ms to 0.25 ms altered the location of ITD functions by 0.04 ms, from 0.07 to 0.11 ms in the example morphology (Fig. 7h, i). Over the population of MSO neuron morphologies, the presence of inhibition reduced firing probability without fundamentally shifting the location of ITD functions (Fig. 7j). However, the region of the function where there is maximal change in firing rate is necessarily impacted by the decrease in ITD width (Figs. 7k, S5), suggesting that dendritic asymmetry and inhibition provide separate contributions to MSO neurons’ receptive field locations.

The simulations in Figs. 4-7 enable one to readily visualize how the shape and timing of EPSPs and IPSPs influence ITD curves. However, *in vivo*, these synaptic currents occur repetitively at varying frequencies and are subject to short term plasticity. To assess the stability of ITD curves under these more complex conditions, we drove the spiking MSO model shown in Figs. 6 and 7 with *in vivo*-like patterns of presynaptic input derived from an auditory nerve model (Zilany et al., 2014), matching total inhibitory and excitatory conductance amplitude and incorporating short-term plasticity measured from MSO neurons in slices (Couchman et al., 2010). IPSP timing was adjusted to lead the EPSPs in accordance with experimental estimates (Roberts et al., 2013). Despite differences in average firing rates of up to 68%, the location and shape of ITDs was generally stable across a range of frequencies in which reliable phase locking is observed (Fig. 8). We did observe a positive shift in the location of ITD functions at 250 Hz. This shift was dependent on the presence of inhibition, but it was not observed at frequencies above 250 Hz (Figs. 8e, S6).

**Fig. 8:**
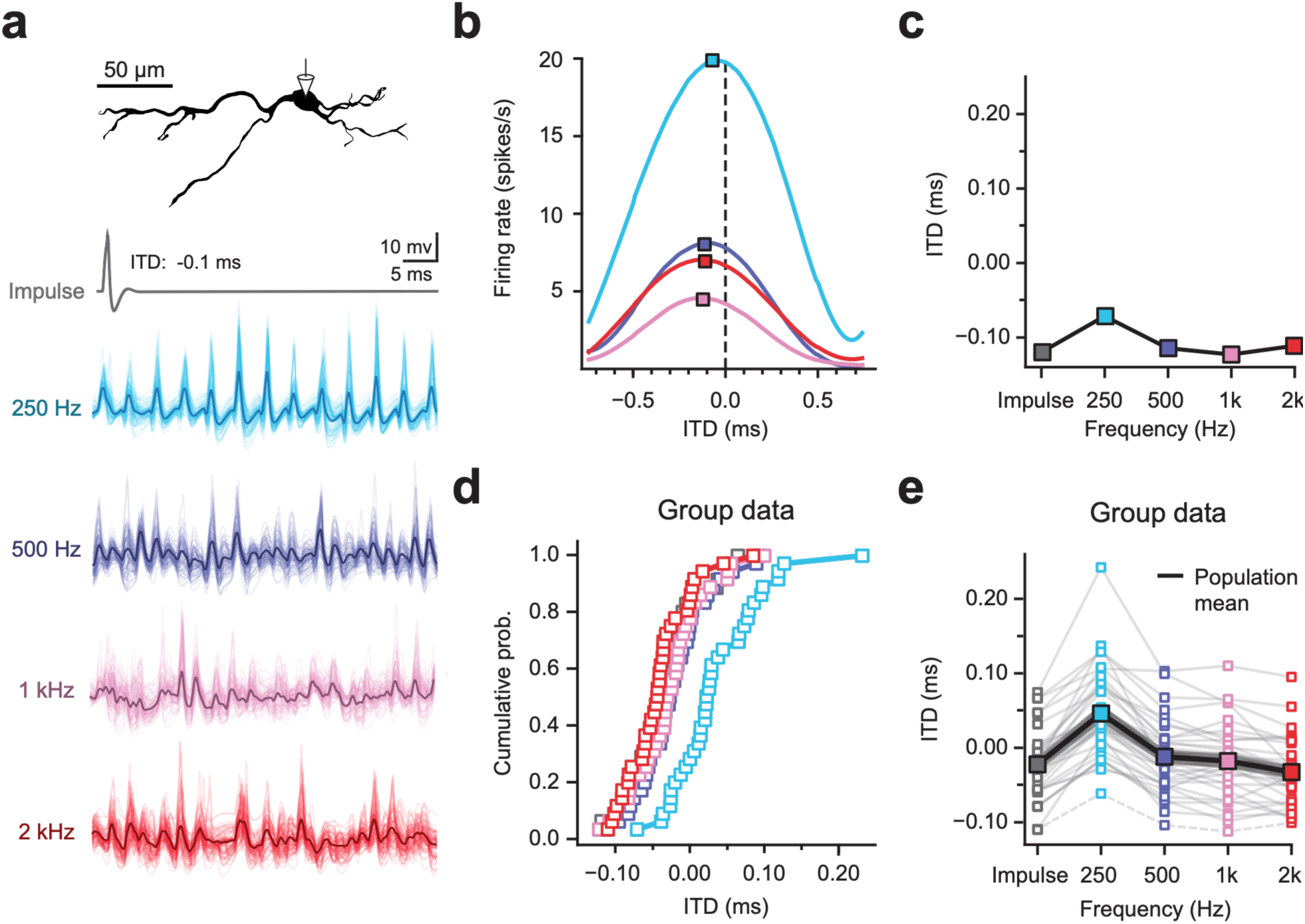
ITD position in responses to synaptic input trains with *in vivo*-like dynamics across frequencies. **a**, Cell morphology (top) and somatic voltage traces across frequencies spanning the phase-locking range (below). Each trace consists of 100 trials (light traces) and an average waveform (dark trace), For comparison, an impulse response is shown to a single synaptic input cycle, as in Figs. 6-7 (top). **b,** ITD tuning curves of the cell in (**a**) across frequencies, with measurements of ITD centroid (squares). Color code as in (**a**). **c**. Centroid vs. stimulus frequency for the example cell in (**a**). Centroids shift at 250 Hz (-0.12 to -0.07 ms), though at higher frequencies they do not (-0.12 to -0.11 ms). **d,** Centroid values across frequency for all 40 cell morphologies. **e**, Centroid values are stable at frequencies above 250 Hz. Gray lines: within-cell measurements. Dashed line: measurements for the cell in **a-c**. Large squares connected by black lines indicate population means.

To explicitly explore the role of branching structure on ITD location, we utilized the model configuration from Fig. 8 and progressively restricted synapse locations to dendrites of decreasing orders (Fig. 9). It can be seen in the example that removal of higher order branches shifts ITD curves rightward, from a slightly ipsilateral input leading curve toward a contralateral leading curve due to the corresponding longer delays on the remaining finer caliber dendrites on the medial side (Fig. 9a-c). Although the direction of shifts across different neurons in our sample set varied according to the details of their dendritic morphology, restriction of innervation to only primary dendrites altered ITD locations by 0.069 ± 0.007 ms on average (Mean ± SEM, p<0.0005, paired Students’ t-test), when compared to uniform innervation of the entire dendritic arbor (Fig. 9d, e).

**Fig 9:**
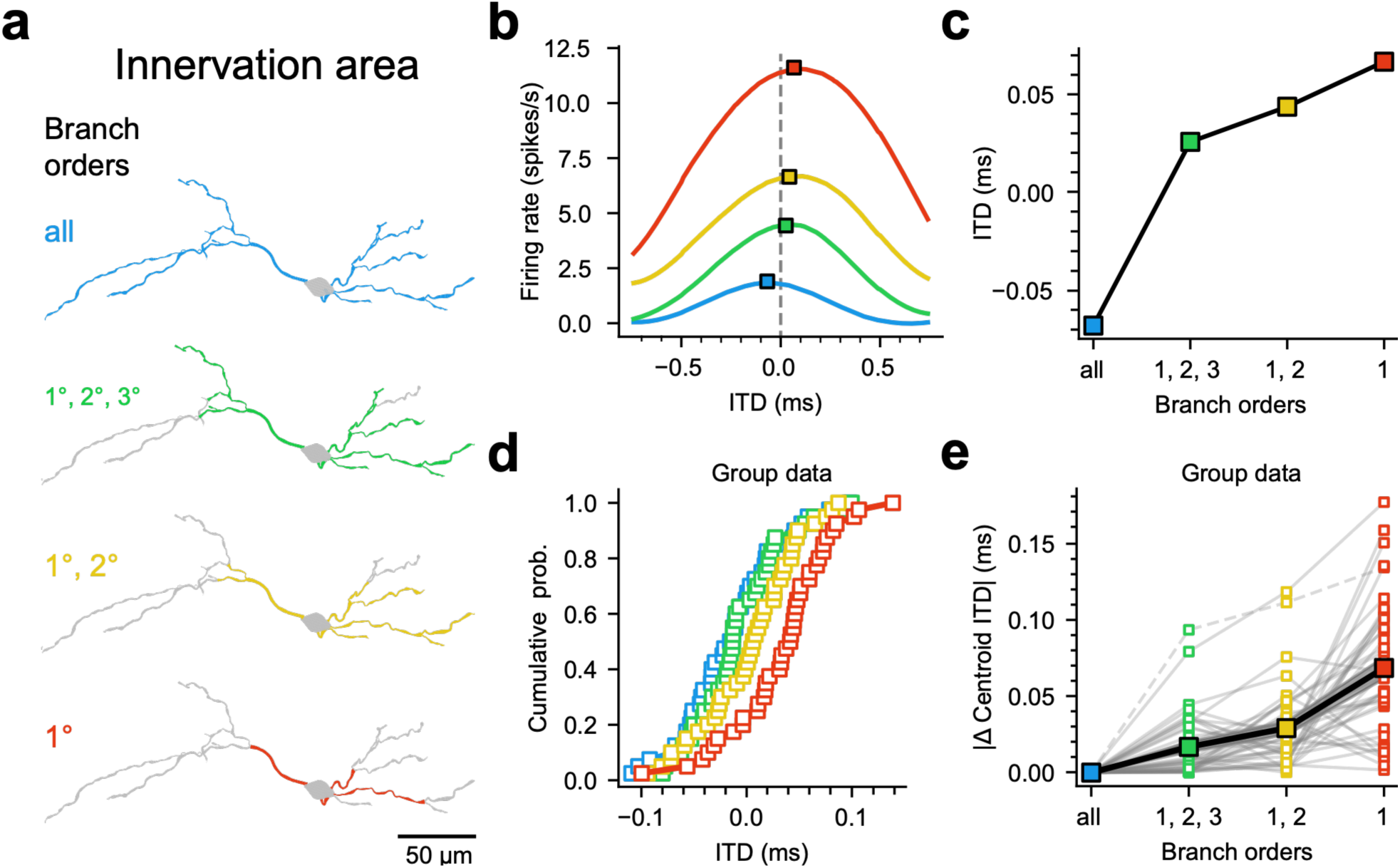
Influence of higher order branches on ITD tuning. a, Branch order restriction of excitatory innervation patterns in an individual neuron for simulation conditions identical to those shown in Fig. 8, at 500 Hz. Branch order color code applies to **b-e**. **b-c,** Normalized ITD tuning curves and measurements of centroid (squares) for each innervation pattern**. d,** Centroid ITDs for the full population of cell morphologies across innervation patterns. **e,** The magnitude of the shift in centroid for each branch order innervation pattern is plotted relative to its difference from the “all” branch order innervation pattern. Cell from **a-c** indicated with a dashed line. The influence of distal synapses to ITD location is reflected in the progressively larger ITD shifts with innervation restrictions to progressively proximal, lower-order branches (green, yellow and red). Larger shifts in individual cells reflect their higher branching complexity.

## DISCUSSION

The passive cable properties of neuronal dendrites are known to impart both attenuation and delay on EPSPs as they propagate to the soma, commensurate with the distance traversed and the anatomical features of the dendrite itself. However, the functional significance of dendritic geometry and its impact on action potential firing is often difficult to assess. Here we have newly identified a critical function for dendritic structure in the processing of sound localization cues in MSO neurons of the auditory brainstem. We have shown that the bipolar dendritic arbor of MSO neurons is typically asymmetrical, imposing a corresponding asymmetry in the relative timing required for excitatory inputs to drive optimal patterns of action potential output. Our compartmental models show that variation in dendritic morphology across cells exerts a continuum of temporal influences sufficient to shift spatial receptive fields across the full biological range of ITDs available to the gerbil. These shifts are insensitive to differences in intensity and likely work together with other sources of internal delay to enable MSO neurons to stably encode spatial locations along the horizontal plane.

### Role of dendritic-based delays in the detection of input sequences

The idea that dendritic structure could dictate detection of temporal sequences of input was first explored in the pioneering computer models of Wilfred Rall. These studies showed that attenuation and propagation delays of EPSPs traveling from the dendrites to the soma could impart greater sensitivity to somatically vs distally directed activation sequences of excitatory inputs, where the longer propagation time of more distal EPSPs is compensated by their earlier activation (Rinzel and Rall, 1974). Preferential firing to centripetal (distal to proximal) temporal sequences of uncaged glutamate at spines has been demonstrated in cortical neurons in slices, but it is not clear what natural conditions would generate such sequences (Branco et al., 2010). By contrast, single excitatory axonal inputs to MSO dendrites mediate a natural centripetal sequence of synaptic activity (Beckius et al., 1999; Callan et al., 2021; Smith et al., 1993), and both current and previous modeling results highlight the importance of matching input conduction velocity with that of MSO dendrites, which conduct EPSPs at an average of ∼0.3 m/s. In the present study, with the aid of 2-photon fluorescence-guided recordings we have been able to sample smaller diameter secondary and tertiary dendrites at longer distances from the soma. Our results reveal that the attenuation and delays of EPSPs propagating from these processes to the soma are more severe than previously estimated (Winters et al., 2017). In our sample of modeled MSO neurons, these slower-conducting non-primary dendrites make up 70% of the surface area of the dendritic arbor in our sample of MSO neurons and thus play an outsized role in exerting temporal shifts in ITD curves in neurons with asymmetrical dendritic trees. These results emphasize that both branching and dendritic diameter are critical determinants of dendritic delay, as seen reflected in the morphoelectrotonic transforms (Figs. 4b,5c). Earlier work reported systematic bilateral asymmetries in excitatory input dynamics and apparent synaptic reversal potentials in MSO neurons (Jercog, 2008; Jercog et al., 2010), which were interpreted as reflecting differences in electrotonic length and synaptic location between medial and lateral dendrites. These functional asymmetries are consistent with the dendritic propagation delays measured here, suggesting that dendritic filtering may underlie previously observed excitatory timing differences.

In the visual system, asymmetrical length and branching of the dendrites drives temporal disparities in the pattern of inhibition of ganglion cells, providing directional tuning to object movement (Bloomfield, 1994, 1991; Trenholm et al., 2011). The computation in the auditory system differs from vision in several fundamental ways. In the retina, spatial information is directly represented at the level of the photoreceptors, and tuning to spatial movement relies on the relative arrival times of excitatory and inhibitory inputs to the dendrites and soma. By contrast, in the cochlea space is not explicitly represented on the basilar membrane, which decomposes complex sounds spatially into different frequency channels. Accordingly, horizontal spatial receptive fields must be constructed *de novo* in the MSO through the detection of synaptic binaural coincidence. In this way, ITDs serve as a proxy for horizontal position.

### Other sources of asymmetry in binaural coincidence detection

Although our data reveal that the dendrites may contribute significant interaural delays in the binaural circuitry in the MSO, they likely work together with other mechanisms that have been proposed to influence binaural coincidence detection. In ∼25% of MSO neurons in gerbils, the axon emerges from either the lateral or medial dendrites (Rautenberg et al., 2009; Scott et al., 2005), and these results appear to extend to other mammalian species, including guinea pigs and cats (Cajal., 1907; Smith, 1995). Results from computer models suggest these offset axons, like asymmetrical dendrites, introduce strong asymmetries in synaptic integration and ITD tuning (Zhou et al., 2005).

Inhibition has received considerable attention as a mechanism for ITD tuning based on shifts of tuning curves measured in single-unit recordings *in vivo* upon iontophoretic block of glycinergic synapses (Brand et al., 2002; Pecka et al., 2008). However, these results are controversial, as strong inhibition has not been observed in both juxtacellular and intracellular patch recordings *in vivo* (Franken et al., 2015; Plauska et al., 2016; Roberts et al., 2013), and there have been concerns about iontophoretically applied strychnine having off-target effects that that alter both firing probability and input resistance (Franken et al., 2015). We found that inhibition did not consistently shift the overall location of ITD curves across the functional range of IPSP magnitudes (Figs. 7,9, S6), consistent with pressure-applied strychnine data *in vivo* (Franken et al., 2015). Inhibition shifted the location of maximal change in firing rate, consistent with a conventional “iceberg” model of inhibitory action on neuronal receptive fields. In mammals, the maximum rate of change, or slope of ITDs may be more important for population encoding of ITDs. Given that the maximal rising slope of firing invariably shifts toward 0 µs ITD when inhibition narrows ITD curves, both inhibition and dendritic asymmetry would both play key roles in setting the location of individual MSO neurons’ receptive fields.

### Comparison with avian binaural coincidence detectors

Binaural coincidence detection has been extensively studied in birds, and thus the comparison to mammals is noteworthy. Unlike mammals, dendritic morphology in bird MSO (nucleus laminaris) shows striking dependence on tonotopic location, exhibiting longer, more highly branched dendrites in neurons from low frequency regions (Smith and Rubel, 1979; Wang and Rubel, 2008). While these tonotopic differences in dendritic length have garnered considerable attention, a close examination of dendritic parameters shows that nucleus laminaris neurons, like those in mammalian MSO, exhibit a continuum of dendritic asymmetry in the length of dendrites receiving ipsilateral and contralateral inputs (Smith and Rubel 1979, their Fig. 15). In these low frequency regions, inputs are restricted to distal, higher order branches (Yamada and Kuba, 2021). Based on both our electrophysiological and modeling data, these higher order branches incur the largest delays as compared to the large-caliber primary dendrites and thus would be the primary determinants of dendritic propagation delay on each side. In the more highly specialized barn owls, the dendrites of laminaris neurons are more limited, and thus internal delay compensation must be mediated primarily by unmyelinated afferents (Carr and Konishi, 1990).

### Dendritic morphology and the scaling problem across species

The biological range of ITDs varies with head size (Jones et al., 2011; Koka et al., 2011), and thus any mechanism of internal delay must be scalable across species, accounting for ITDs differing by hundreds of microseconds (e.g., gerbil: ±130 µs (Maier and Klump, 2006), human: ± 600-700 µs (George, 1977)). A dendrite-based source of internal delay would be easily scalable, as the length of MSO dendrites increases with the size of different species. Although quantitative data are not available, previous morphological studies in guinea pig, ferret, cat and humans show that the dendritic arbors show considerable asymmetries (Cajal., 1907; Henkel and Brunso-Bechtold, 1990; Kulesza, 2008; Smith, 1995). Thus, the functional impact of dendritic asymmetry we describe in gerbils may well reflect a structural phenomenon found more broadly across mammalian species.

## METHODS

All procedures were conducted in agreement with the Institutional Animal Care and Use Committee at The University of Texas at Austin (UT-Austin), which followed the guidelines of the National Institutes of Health. Mongolian gerbils (*Meriones unguiculatus*) were housed and raised in a colony at the UT-Austin Animal Resource Center. Animals experienced a 50/50 day/night cycle and had continuous access to food and water.

### Brain slice preparation

both male and female gerbils (17-40 days old) were anesthetized with isoflurane and decapitated upon loss of the choroid reflex. The brain was removed from the cranium under warm (34-35°C) artificial cerebrospinal fluid (ACSF), the cerebellum was removed and the isolated hindbrain was glued to the stage of an oscillating tissue slicer (Leica VT-1200S). Horizontal sections were prepared at 200 µm thickness and stored at 35°C in a holding chamber for 45-60 min., and then gradually cooled to room temperature (24°C). All solutions were continuously bubbled with 95% O_2_ / 5% CO_2_. ACSF contained (in mM): 125 mM NaCl, 25 mM D-glucose, 2.5 mM KCl, 25 mM NaHCO_3_, 1.25 mM NaH_2_PO_4_, 1.5mM MgSO_4_, 1.5 mM CaCl_2_, pH adjusted to 7.45 with NaOH, and final osmolarity at 315 mOsm.

### Imaging and electrophysiological acquisition and analyses: neurons in the MSO were

visualized under gradient contrast optics on a Leica TCS SP5 System equipped with a Coherent Chameleon Ultra II Ti:sapphire laser and an 8 kHz resonant scanner under control of Leica LAS AF Imaging Software. We found that gradient contrast images acquired via the photomultiplier tubes were of poor contrast in the heavily myelinated regions of the auditory brainstem, and so cell selection and seal formation was carried out under gradient contrast imaging through the epifluorescence port of the microscope and detected on a high-resolution monochrome digital camera (Zeiss Axiocam 503) running Zen software. The soma was recorded in whole-cell current clamp mode with a pipette (4 to 8 MΩ resistance) containing a potassium gluconate-based solution with 40-80 µM Alexa 594 added for visualizing cell morphology. After waiting ∼10 min. for the dye to fill the dendritic arbor, a dendritic recording was made under 2-photon fluorescence with a second pipette (8 to 15 MΩ resistance) containing the same K Gluconate solution with 40-80 µM Alexa 488. Pipette solutions had a composition of 115 mM K-gluconate, 4.42 mM KCl, 0.5 mM EGTA, 10 mM HEPES, 10 mM, Na_2_Phosphocreatine, 4 mM MgATP, and 0.3 mM NaGTP. Osmolality was adjusted to 300 mOsm/L with sucrose, and pH was adjusted to 7.30 with KOH. Seal resistances exceeded 1 GΩ. Series resistance and capacitance compensation were implemented, and recordings were discontinued if series resistances exceeded 20 or 50 MΩ for somatic and dendritic recordings, respectively. Membrane time constants were measured from small (3-5 mV) hyperpolarizing voltage responses to current injections in the somatic or dendritic recording pipettes. Electrophysiological recordings were under the control of custom routines written in IGOR-Pro (Wavemetrics). Pipettes were prepared on a Sutter P-1000 puller using borosilicate glass (1.5 mm O.D.). Data was acquired using a pair of Dagan BVC-700, amplifiers in conjunction with an ITC-18 computer interface (Heka Instruments). Data was low-pass filtered at 5 kHz, sampled at 50 kHz, and stored on a Macintosh computer for further analyses using custom-written routines in IGOR-Pro.

### Compartment model construction

A subpopulation of 40 fully intact reconstructed MSO neurons was selected from those gathered in Bondy et al., 2021. These neurons were chosen exclusively on the basis of their morphological integrity and consisted of primarily phasic firing neurons (38 of 40 cells). The remaining two neurons exhibited an “oscillating” pattern of repetitive firing. The average membrane time constant of these neurons during 3-5 mV hyperpolarizations was 0.31 ± 0.02 ms, and their ages ranged from 19-21 postnatal days. Compartmental models were uniformly applied to all morphologies, and we did not attempt to mimic each neuron’s individual properties. The reconstructed morphologies were imported into the NEURON simulation environment, using its Neurolucida importing tool. Computational compartments were created every 2 μm (Fig. S7a). Channels were implemented into the model, consisting of low and high-voltage-activated potassium channels, hyperpolarization-activated, cyclic nucleotide-gated (HCN) channels, leak channels, and sodium channels in accordance with densities and distributions in previous observations (Table 1; Mathews et al., 2010; Nabel et al., 2019; Khurana et al., 2012; Scott et al., 2010). In simulations modeling a passive cell, high and low-voltage potassium and HCN channels were replaced with passive leak channels with conductances set to the average resting conductance of the preexisting voltage-gated channels. In simulations with an axon, a model axon with morphology as in Lehnert et al., 2014 was attached at the center of the soma. The axon channel kinetics implemented consisted of sodium channels, low-threshold potassium channels and leak channels in the nodes and initial segments, with solely leak channels in the internodes. Measurements from the axon were all recorded from the fifth node. The kinetics of synaptic conductances were described by an alpha function (τ = 290 µs).

**Table 1:**
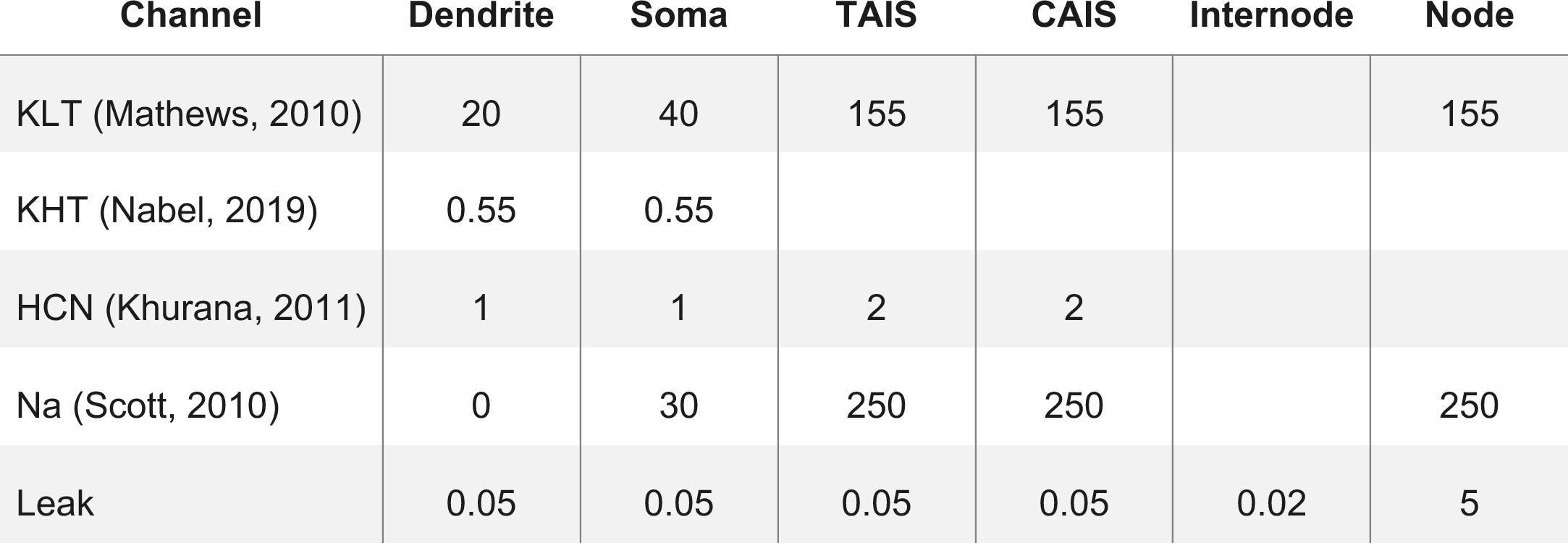
Maximal channel conductances (µS/cm^2^). Tapered axon initial segment (TAIS) and constant diameter axon initial segment (CAIS).

| Channel | Dendrite | Soma | TAIS | CAIS | Internode | Node |
| --- | --- | --- | --- | --- | --- | --- |
| KLT (Mathews, 2010) | 20 | 40 | 155 | 155 |  | 155 |
| KHT (Nabel, 2019) | 0.55 | 0.55 |  |  |  |  |
| HCN (Khurana, 2011) | 1 | 1 | 2 | 2 |  |  |
| Na (Scott, 2010) | 0 | 30 | 250 | 250 |  | 250 |
| Leak | 0.05 | 0.05 | 0.05 | 0.05 | 0.02 | 5 |

### Simulations

When measuring EPSP propagation, synapses were placed every 2 µm, and unless otherwise noted they were given a unitary conductance that would maximally depolarize the soma 1 mV. In simulations of ITD coincidence detection, dendrites were innervated with synapses (release probability=0.45) placed every 2 μm, totaling ∼125 synapses for each side (Couchman et al., 2010). To mimic sequential synaptic release along inputs, an artificial axonal input delay (Δt_axon_) was calculated for each synapse as a function of conduction velocity (ν_input_) and axonal travel distance. The travel distance was considered zero at each dendritic arbor’s furthest point from the soma. Otherwise, the axonal travel distance consists of two components: the mediolateral distance from the furthest point (Δt_axon_=0) to the synapse’s closest terminal segment; the path length from that terminal segment to the synapse (Fig. S7b). Simulated acoustic ITDs could then be added to the internal calculated axonal delay. Each simulated acoustic ITD value (-0.5 to 0.5 ms, incremented by 0.01), was tested with 400 trials, and each trial consisted of a single stimulation of all synapses. To provide a more binary output, a model axon was attached to the cell’s soma, and a spike was reported if a 25 mV depolarization from rest was recorded at the fifth and final axonal node. The synapses could then be adjusted with a unitary conductance to achieve an acceptable maximum firing probability. When implementing inhibition in coincidence detection simulations, two inhibitory synapses were added at the soma, with each respectively driven from ipsilateral and contralateral sides. Each synapse generated an IPSP whose kinetics were described by an alpha function with a time constant of 1.5 ms (Couchman et al., 2010). The timing of the synapses was adjusted relative to the artificial excitatory ITD signals. The amplitude of the IPSPs was adjusted by modifying the inhibitory synapses’ conductances to achieve a combined IPSP hyperpolarization of approximately 1, 2, and 3 mV.

A realistic spiking auditory nerve fiber (ANF) model was used to generate synaptic input trains (Zilany et al., 2014). Spike trains from high spontaneous rate fibers were generated using a sinusoidal input at a given frequency, which was also used as the characteristic frequency of the model fiber. To accommodate the new input type, stochastic synapses exhibiting synaptic depression were implemented as in Callan (2021), with a model vesicle pool of 40 for each synapse. Each arbor was innervated by axonal inputs equal to the number of terminal branches, and each input was assigned a random pre-generated spike train. The same process was done for the two inhibitory inputs. Total excitatory and inhibitory conductance values were matched. Each stimulus was presented for a duration of 200 ms and the range of simulated acoustic ITDs was increased to ± 0.75 ms. In simulations regarding branch order, we selectively innervated subsets of branches based on their order. The three innervated subsets consisted of primary, primary and secondary, and primary through tertiary. These selective innervation experiments were performed with an ANF input generated by a 500 Hz simulated sound stimulus.

#### Data analyses

Measures of dendritic distance, both in slice experiments and in compartment models, were taken as the length along the dendritic arbor to the center of the somatic compartment. To address the variable size of the MSO within the slice plane, a normalized tonotopic location was calculated as the cell’s distance from the most dorsal point of the MSO, relative to the entire MSO dorsoventral length. To provide a quantitative value of asymmetry between each cell’s morphological measures, a unit of fold difference was used, with a value of 1 and 2 representing equal and double values, respectively. Average path length measurements were taken from the distances along the dendrite to the soma from each terminal section of the corresponding mediolateral arbor. Measurements for propagation attenuation and timing were taken at both the synaptic and somatic compartments of the model cell. For attenuation, the somatic depolarization for mediolateral sides was calculated by averaging each synaptic compartments propagated EPSP amplitude at the soma. Morpho-electrotonic transforms were created by scaling each sections length according to the natural logarithm of its attenuation 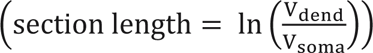. EPSP delay values were calculated by measuring the time between peak depolarization in the synaptic and somatic compartments. The corresponding delay transforms are scaled according to the delay contribution of each section. Mediolateral comparisons of delay were calculated by subtracting each arbor’s average EPSP delay for all compartments. To minimize noise and data filtering for the ITD response, the centroid of the curve was measured, where half of the total area under the curve exists on either side. Comparisons were made using the absolute ITD centroid values (Figs. 6e-f,9e) to highlight the population of neurons’ collective change. The ITD firing probabilities were smoothed with a Savitzky–Golay filter for the purpose of reasonably identifying the region of maximum slope for the curves.

