## Supplementary figures for "Dendritic delay lines shape the computation of sound location in neurons of the gerbil medial superior olive"

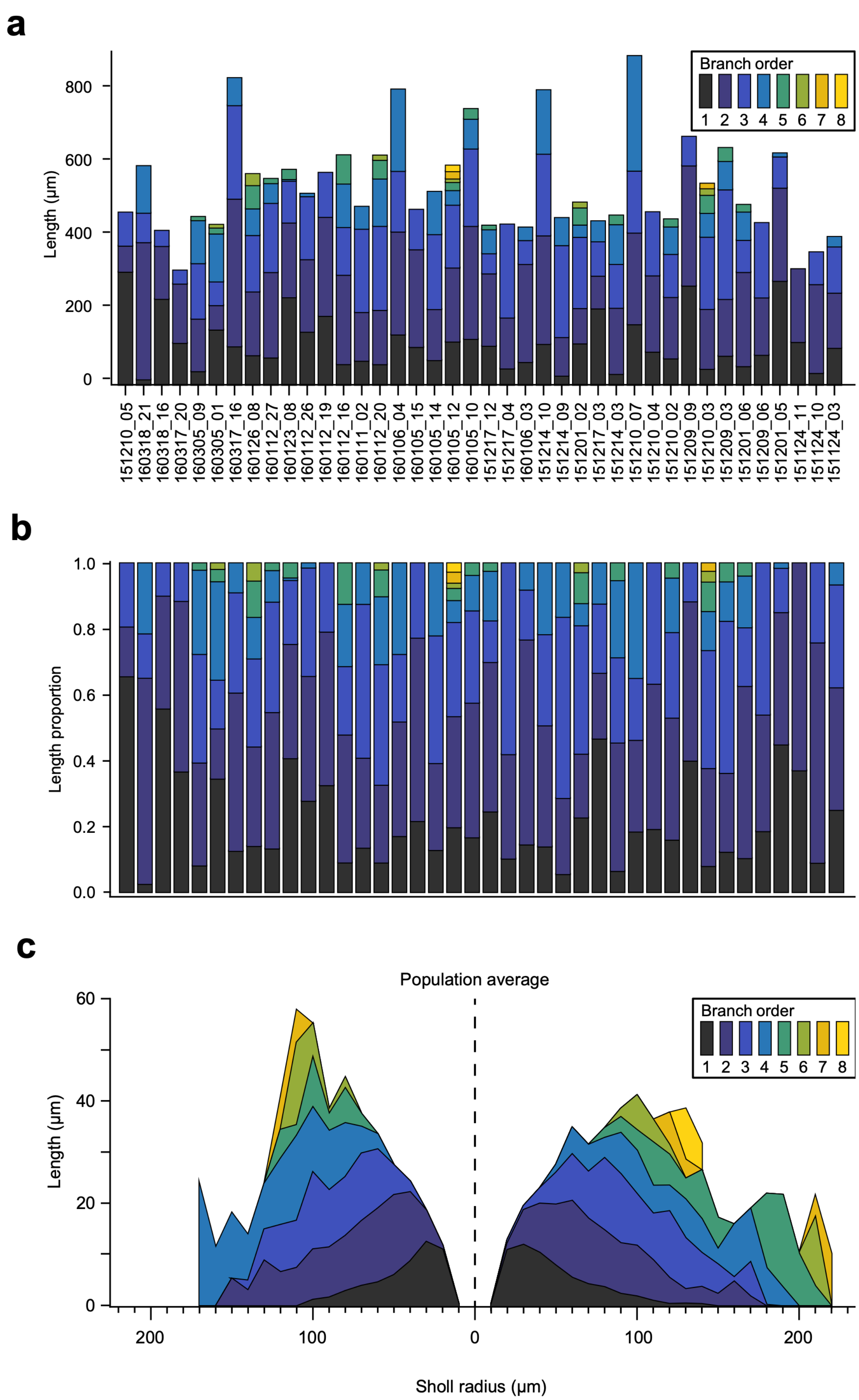
a
Branch order
1 2 3 4 5 6 7 8
Length (µm)
0
200
400
600
800
151210_05
160318_21
160318_16
160317_20
160305_09
160305_01
160317_16
160126_08
160112_27
160123_08
160112_26
160112_19
160112_16
160111_02
160112_20
160106_04
160105_15
160105_14
160105_12
160105_10
151217_12
151217_04
160106_03
151214_10
151214_09
151201_02
151217_03
151214_03
151210_07
151210_04
151210_02
151209_09
151210_03
151209_03
151201_06
151209_06
151201_05
151124_11
151124_10
151124_03
b
Length proportion
0.0
0.2
0.4
0.6
0.8
1.0
c
Population average
Branch order
1 2 3 4 5 6 7 8
Length (µm)
0
20
40
60
200
100
0
100
200
Sholl radius (µm)

**Figure S1: Measurements of branching in morphology population.** A: Absolute (left) and relative (right) length of all 40 reconstructed dendritic arbors, colored according to the degree of each branch, with 1º being the primary branches. Average proportions of dendritic arbors for the population (± SD): 1º: 0.2988 ± 0.15, 2º: 0.3702 ± 0.12, 3º: 0.2297 ± 0.11, 4º: 0.0827 ± 0.08, 5º: 0.0149 ± 0.03, 6º: 0.0026 ± 0.01

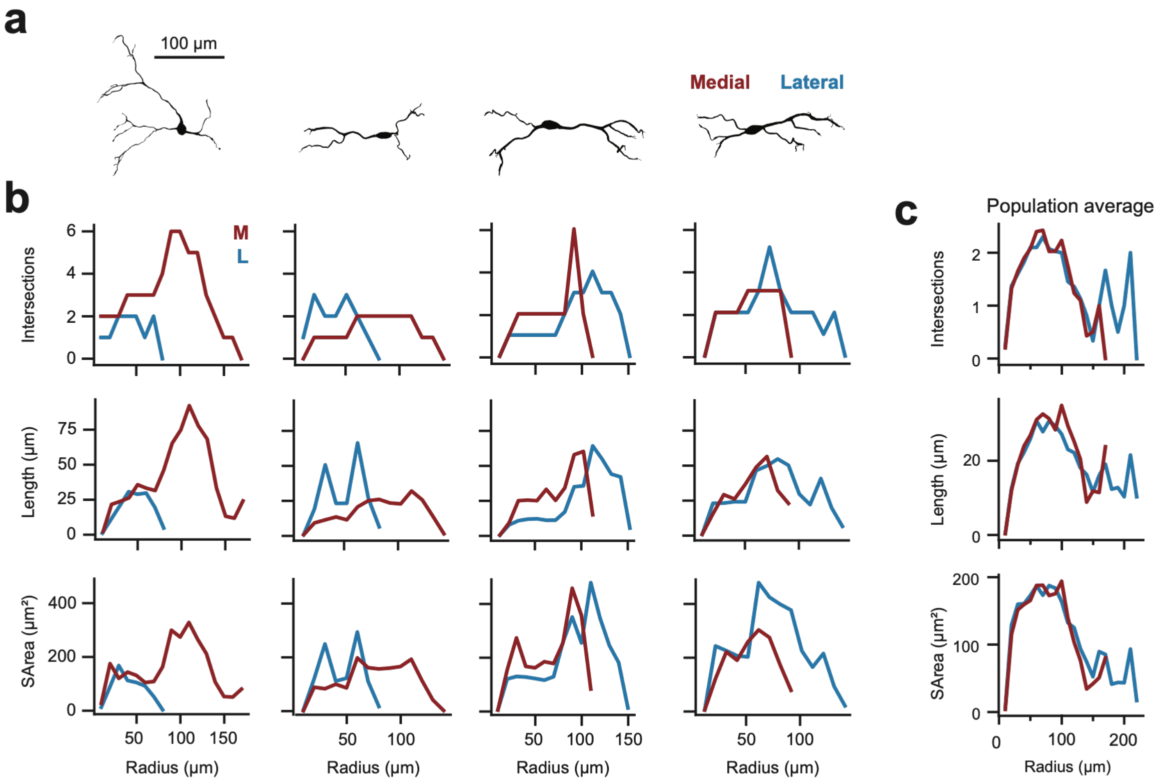


**Figure S2: Averaging dendritic measurements obfuscates cell diversity. a,** Four example cell morphologies used in **b**, below**.** Medial (red, left) and lateral (blue, right) is indicated. **b,** Sholl profiles for each mediolateral side across three measures of dendritic morphology (intersections, path length, and surface area). Colors correspond to **a**. Individual cells demonstrate unique profiles. **c,** Sholl profiles (as in **b**) averaged across the population. Averaged mediolateral profiles show qualitatively small differences, unlike individuals in **b.**

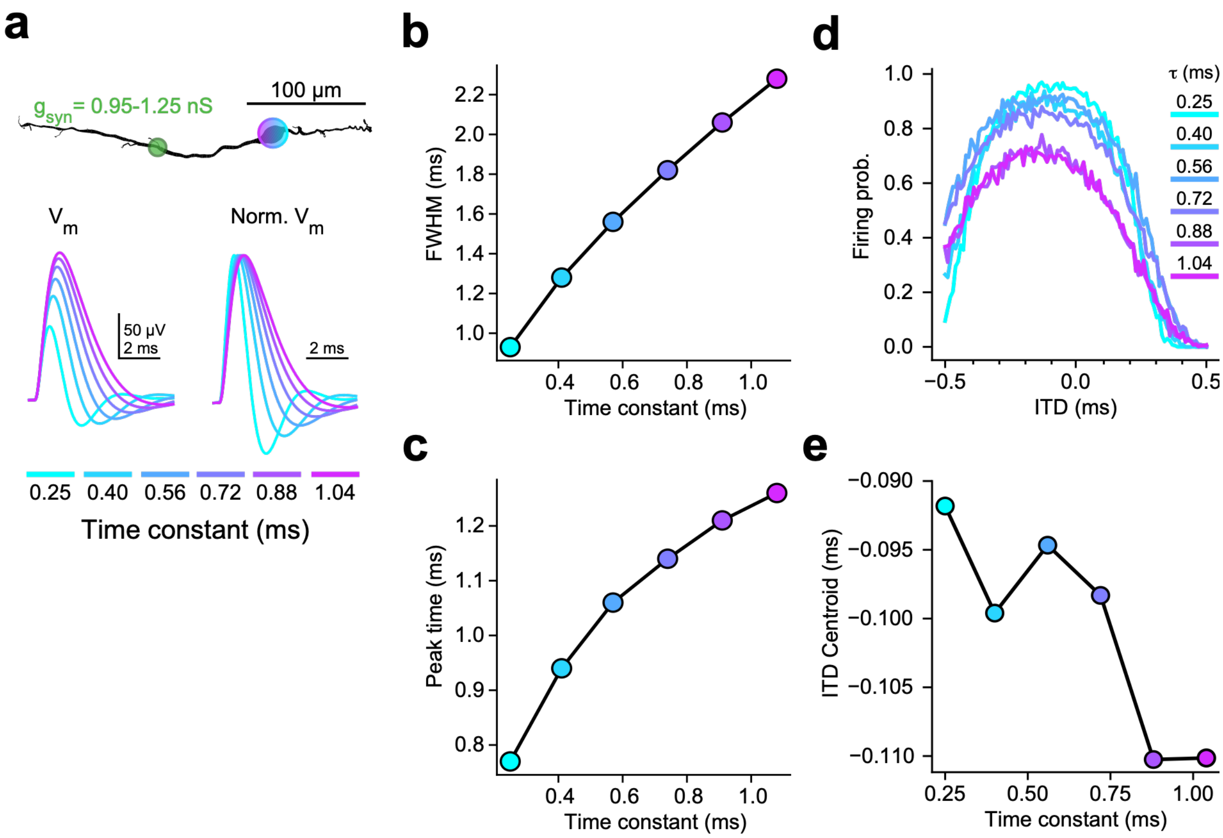


**Figure S3: ITD tuning minimally sensitive to model physiology. a,** Exemplar cell morphology (top) with a marking at the soma (gradient), where traces (bottom) were measured, and a marking at the site of synaptic input (green, maximal conductance listed below). Nonnormalized (left) and normalized (right) and EPSP traces driven by synaptic input, for each time constant adjustment. Time constant measurements are indicated by the legend below. **b,c,** Full width half maximum (**b**) and peak time (**c**) measurements of EPSP traces from **a**, against its manipulated time constant measurement. Colors corresponding to A, for consistency. **d,** ITD tuning curves for each tested time constant, colored according to the inlayed legend (same as **a-c**). E: Centroid measurements (0.092 – 0.110 ms) for each ITD tuning curve from **d**, against its manipulated time constant.

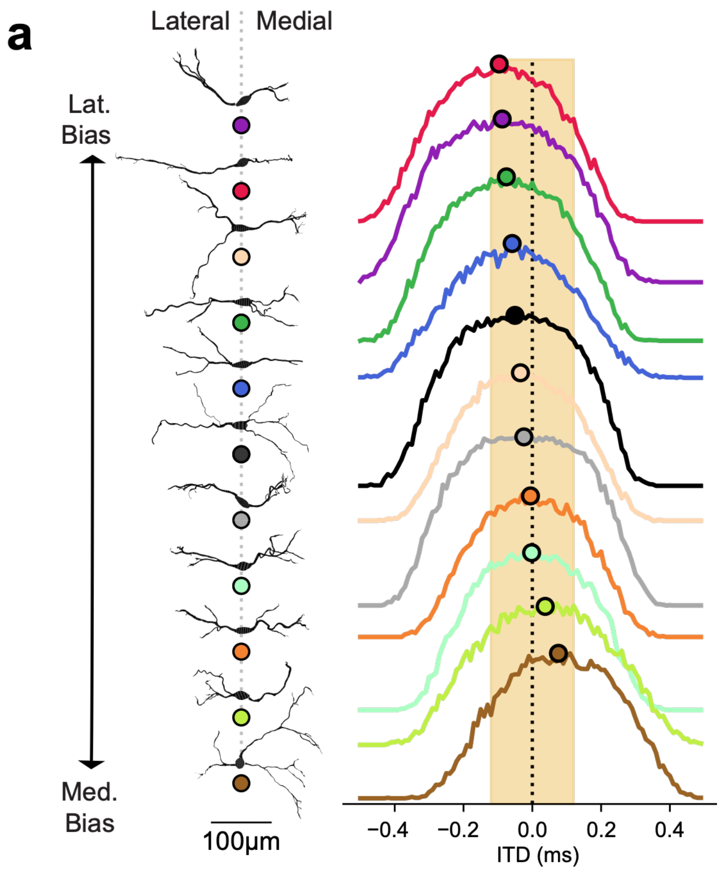


**Figure S4: Morphologies visually correspond to ITD tuning curves. a,** Reconstructed morphologies (left), sorted according to average path length, as in Figure 3A. ITD tuning curves (right) sorted according to curve centroids (-0.096 to 0.076 ms), offset vertically from each other. Curves colored and centroids marked with corresponding color from sorted morphology. Same coloring as Figure 3. Ecologically possible ITD range of model species indicated in light orange. Dotted line indicates at ITD = 0 ms. Three cells excluded due to simulation constraints.

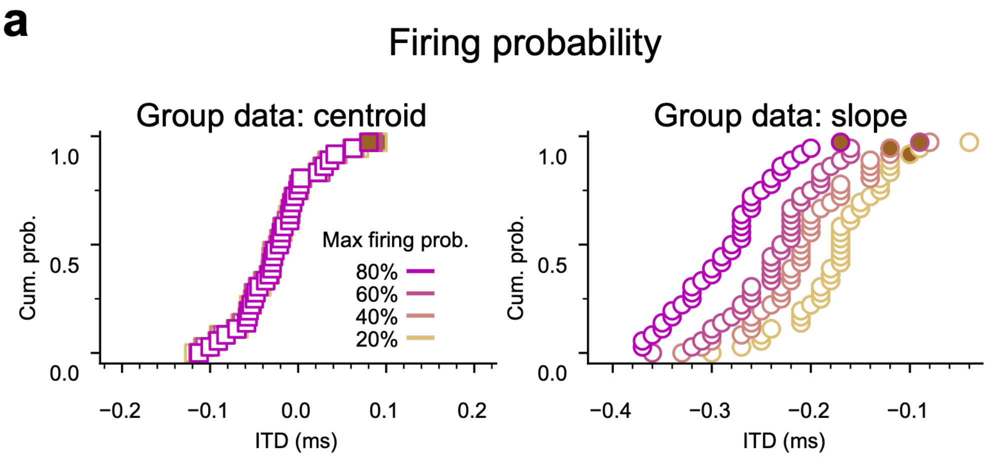


**Figure S5: Input intensity produces shifts in tuning curves' rising edges. a,** Group data (n=36) for ITD centroids and point of maximal slope for each firing probability, representing stimulus intensity. Magnitude of depolarization reproduces the shift of tuning curves maximal slopes in the case of inhibition, as seen in Figure 7F.

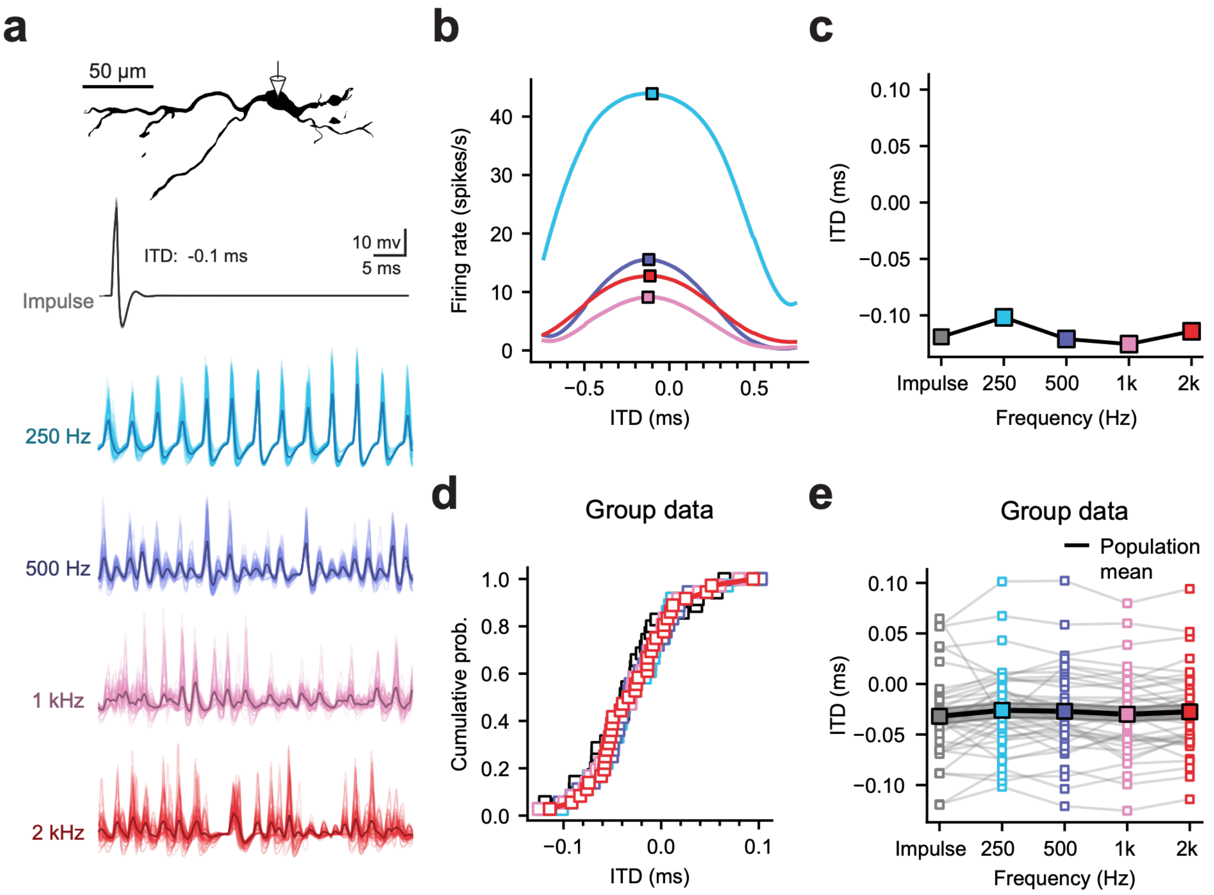


**Figure S6: ITD position in responses to synaptic input trains with *in vivo*-like dynamics across frequencies without inhibition.** **a**, Cell morphology (top) and somatic voltage traces across frequencies spanning the phase-locking range (below). Each trace consists of 100 trials (light traces) and an average waveform (dark trace), For comparison, an impulse response is shown to a single synaptic input cycle, as in Fig. 6-7 (top). **b,** ITD tuning curves of the cell in (**a**) across frequencies, with measurements of ITD centroid (squares). Color code as in (**a**). **c**. Centroid vs. stimulus frequency for the example cell in (**a**). Centroid shift at 250 Hz was not observed, compared to shift in Fig 8c with inhibition. **d,** Centroid values across frequency for all 40 cell morphologies. **e**, Centroid values are stable at frequencies above 250 Hz. Gray lines: within-cell measurements. Dashed line: morphology in **a-c**. Large squares connected by black lines indicate population means.

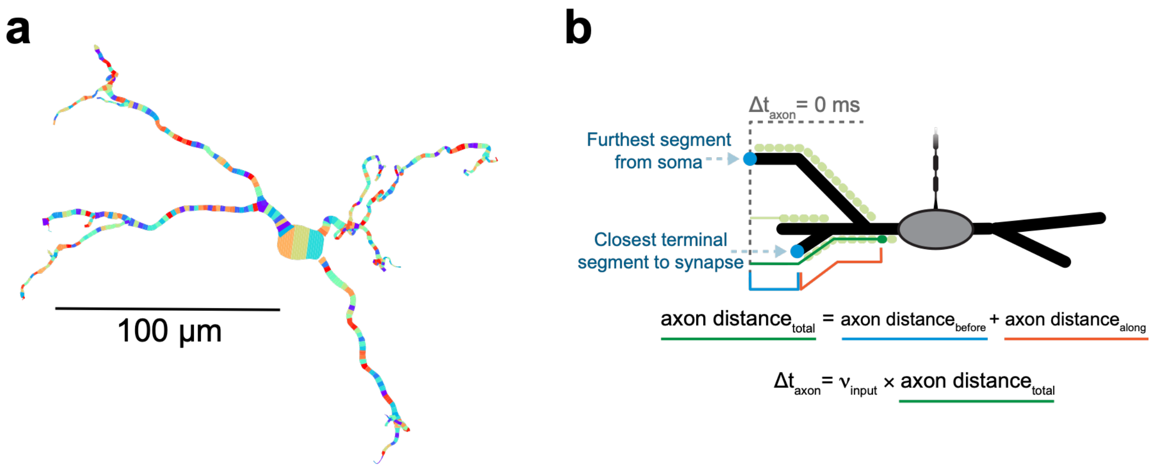


**Figure S7: Modeling visual aids. a,** Morphology of one reconstructed MSO neuron, with colored patterning to distinguish each 2 μm computational compartments created. **b,** Cartoon illustrating artificial axonal input definition for a mediolateral side. Grey dotted line indicates point at which there is no axonal input delay to a synapse. Synapse used for demonstration in dark green, with others not demonstrated in light green. Total axonal travel distance (green) consists of two parts: mediolateral distance from side's furthest segment and the closest terminal segment to the synapse (blue); path length from the closest terminal segment to the synapse (orange).
